# Blockade of mTOR ameliorates IgA nephropathy by correcting CD89 and CD71 dysfunctions in humanized mice

**DOI:** 10.1101/2025.01.21.634071

**Authors:** Alexandra Cambier, Jennifer Da Silva, Julie Bex, Fanny Canesi, Lison Lachize, Aurélie Sannier, Natacha Patey, Erwan Boedec, Amandine Badie, Renato C. Monteiro

**Author notes:** in memorium (Natacha Patey).

## Abstract

**Background and hypothesis:** IgA nephropathy (IgAN) is the most common glomerulonephritis and major cause of renal failure worldwide. Pathogenesis involves galactose deficient IgA1-immune complexes containing soluble IgA Fc receptor (sCD89). Both IgA1 and sCD89 bind independently to transferrin receptor (TfR1, CD71), a mesangial IgA1 receptor. Here, we hypothesize that sCD89 plays a pathogenic role in IgAN by driving a tri-partite IgA1-sCD89-CD71 complex inducing activation *in situ* of the mTOR (mammalian target of rapamycin) signaling pathway in mesangial cells and contributing to disease progression. mTOR inhibition may disrupt this pathogenic axis, reducing IgA1 and sCD89 deposits, modulating CD71 expression, and alleviating disease manifestations. Here, we investigated the role of sCD89 and a mTOR inhibitor using humanized mouse models of IgAN expressing CD89 and/or IgA1.

**Methods:** Single cell and RNAseq were obtained from IgAN public dataset and immunostaining from childhood IgAN (cIgAN) biopsies. Human mesangial cell (HMC) stimulation by recombinant sCD89 (rsCD89) was followed by western blot analysis. Pre-clinical assays with mTOR inhibitor (Everolimus) by oral gavage were performed using young α1^KI^ mice injected with rsCD89 for 25 days (preventive protocol) and adult α1^KI^CD89^Tg^ mice (treated protocol) for 75 days. Proteinuria, renal function, and circulating immune-complexes (CIC) were analysed and kidneys harvested for histology. Results: RNAseq revealed increased TfR1 and mTOR mesangial cell expression in IgAN patients. TfR1 upregulation was confirmed in cIgAN biopsies. sCD89 stimulation induced HMC TfR1 expression and phosphorylation of mTOR, Akt and p70S6K1. Everolimus treatment prevented or reverted mesangial IgA1 and C3 deposits but also decreased mesangial TfR1 and cell proliferation. Everolimus impaired levels of sCD89- and IgA-CIC, proteinuria, as well as renal function.

**Conclusion:** These findings highlight the critical role of the sCD89-TfR1-mTOR axis in IgAN pathogenesis and support the use of mTOR inhibitors as a novel therapeutic approach. This approach could significantly improve outcomes by slowing disease progression and minimizing the systemic toxic effects of current immunosuppressive therapies. This is particularly crucial for pediatric patients, where the only approved treatment – steroids – has severe side effects, including detrimental impacts on bone health and growth.

## Introduction

IgA nephropathy (IgAN) is an autoimmune glomerular disease associated with environmental factors and genetic polymorphisms leading to a significant health burden (1, 2). It usually progresses over 20 years to end-stage renal disease (ESRD) in one third of patients (3-5), requiring kidney transplantation early in life (6-9).

IgAN pathogenesis presents four consecutive hits: (1) generation of abnormal galactose-deficient IgA1 (Gd-IgA1)(10, 11), (2) an auto-immune reaction that generates IgG autoantibodies targeting the Gd-IgA1, (3) formation of circulating IgA1 immune complexes (CICs) composed of IgG anti-Gd-IgA1 and other components (12). Gd-IgA1 triggers the myeloid IgA Fc receptor CD89 leading to its shedding from the cell surface (13-15). Soluble forms of CD89 (sCD89) generate and amplify the size of CICs (14) forming larger Gd-IgA1-sCD89 CICs(16-21). (4) The Gd-IgA1-sCD89 CIC specifically deposits in the glomerular mesangium and, through at least partially by interactions with the transferrin receptor 1 (TfR1 also known as CD71), a mesangial IgA1 receptor, activating the complement cascade and glomerular inflammatory response (22). CIC activation of human mesangial cells (HMCs) enhances cell proliferation with mesangial matrix expansion and pro-inflammatory cytokine secretion. In turn, HMC activation induces podocyte hypertrophy or sclerosis leading to proteinuria and hematuria and thus the progression of IgAN (23-25).

Mesangium is one of the main sites for immune-complex deposition resulting in cell proliferation and increased extracellular matrix production during glomerulonephritis development. In IgAN, mechanisms of proliferation of human mesangial cells (HMCs) are poorly understood. TfR1 has been described as an IgA1 receptor in IgAN but also in celiac disease (26, 27). TfR1 is overexpressed in the mesangium of IgAN patients, and it is associated with IgA deposits (28). Deglycosylation of IgA1 increases the TfR1-IgA interaction (29) and HMC stimulation by polymeric IgA1 enhances TfR1 expression on these cells leading to a positive feedback loop favoring IgA1 glomerular deposition (30).

The mechanism by which IgA1 induces TfR1 overexpression in mesangial cells remains poorly understood. It has been demonstrated that mesangial TfR1 overexpression in IgAN is influenced not only by Gd-IgA1 but also by sCD89 and transglutaminase 2. Among these, sCD89 plays a pivotal role in the selective deposition of IgA1 in the mesangium, as evidenced by studies in a humanized mouse model expressing IgA1 either alone or in conjunction with CD89 (21, 31). In a recent study, we identified various forms of soluble CD89 in the serum of patients with childhood IgAN (cIgAN), with or without its association with IgA1 deposits in the mesangium (21). Patient-derived and recombinant sCD89 forms significantly activate HMCs and induce their proliferative state through interaction with TfR1 (21). This interaction activates the PI3K/Akt/mTOR signaling pathway, which is associated with a poor prognosis in IgAN (21, 31, 32).

The PI3K/Akt/mTOR signaling pathway accelerates the cell life cycle, reduces apoptosis, and promotes cell migration (33, 34). mTOR inhibitors can represent an attractive therapeutic option since oral corticosteroids is the only approved immunosuppressor that has considerable systemic toxic effects, especially in pediatrics (35). In this study, we hypothesize that sCD89 directly interacts with TfR1 to activate the PI3K/Akt/mTOR signaling pathway, driving mesangial cell proliferation and contributing to IgAN progression. Inhibiting mTOR could disrupt this pathogenic axis, reducing mesangial hyperplasia, immune complex deposition, and disease severity, thereby offering a promising therapeutic strategy for IgAN.

## Methods

### Mouse procedures

Mice expressing the IgA1 heavy chain (α1^KI^) and transgenic for the human Fc α receptor I (CD89^tg^), which spontaneously develop IgAN at 12 weeks of age as previously described (α1^KI^CD89^tg^ mice (31)), were used as indicated. We also utilized α1^KI^ mice expressing human IgA1 (36), which were injected with recombinant sCD89 (rsCD89) over a 4-week period. Mice were weaned at 4 weeks of age, housed under specific pathogen–free (SPF) conditions, provided sterile water to drink (supplemented as indicated), and fed sterilized chow ad libitum. Ethical approval for animal experiments was granted by the local animal ethics committee (Autorisation de Projet Utilisant des Animaux à des Fins Scientifiques, APAFIS no. 42753). Mice were anaesthetized before terminal collection of blood from the retro-orbital sinus and then euthanized by cervical dislocation. Euthanized mice were perfused with PBS. Segments of kidney were snap-frozen in optimal cutting temperature (OCT) medium using liquid nitrogen. Kidneys were also fixed in 4% formaldehyde. Segments of kidney were snap-frozen dry in cryotubes using liquid nitrogen.

### Experimental procedures

For each protocol, 6 α1^KI^ mice (that received rsCD89 injection) and 20 α1^KI^CD89^Tg^ mice have been used to make two arms: a treated arm with the mTOR inhibitor everolimus at standard dose (2 mg/kg) and a control arm receiving a placebo treatment with water by daily intragastric gavage for each mouse model. The dosage has been reassessed with the weight of the mice. Everolimus has been dissolved in sterile water before administration by tube feeding. A first blood sample has been taken before any intervention to measure the kidney function of the mice as well as the level of CICs. Urine has been collected to monitor mouse proteinuria before and after treatment. After 25 days for the α1^KI^ (with sCD89 injection) and 75 days for α1^KI^CD89^Tg^, the mice have been sacrificed. Plasma and serum have been collected to measure renal function and CICs. The CICs (IgA-IgG, sCD89-IgA), Gd-IgA1 and sCD89, have been assessed by sandwich ELISA as described before (21). The kidneys were harvested for histological analysis, including: (1) Ki67 staining to evaluate kidney hypercellularity, (2) assessment of human IgA1 and complement C3 deposits by immunofluorescence, and (3) evaluation of transferrin receptor expression (anti-CD71 antibody) and CD89 expression (A3 monoclonal anti-CD89 antibody) by immunohistochemistry. Quantification of immunostainings was performed in 10 glomeruli per mouse at the end of the treatment.

### Production of soluble proteins

The recombinant soluble CD89 (rsCD89) was produced at the Biochemistry and Biophysics (B&B) facility (Paris, France). Briefly, the human sCD89 sequence was codon-optimized (GeneArt, ThermoFisher, USA) to include an N-terminal His tag and TEV cleavage site and then inserted into the pCDNA3.4 vector (ThermoFisher, USA) using the NEBuilder HiFi DNA Assembly Kit (New England Biolabs, USA). The resulting pCDNA3.4-sCD89 plasmid was purified using the NucleoBond Xtra Maxi EF Kit (Macherey-Nagel, Germany) and verified by sequencing (Eurofins Genomics, Germany). The pCDNA3.4-sCD89 plasmid was transfected into Expi293 cells (ThermoFisher, USA) at a density of 3 × 10^6^ cells/mL using ExpiFectamine (ThermoFisher, USA). The cells were incubated for 6 days at 37°C, 5% CO<sub>2</sub>, and 125 rpm. After incubation, the supernatant containing rsCD89 was centrifuged at 6,000 × g for 15 minutes and purified using a HisTrap Excel column (Cytiva, Sweden) on an ÄKTA Pure chromatography system (Cytiva, Sweden). Elution was performed with an imidazole gradient, and the rsCD89 was desalted into PBS 1X using a HiPrep 26/10 desalting column (Cytiva, Sweden).

### Cell culture and treatments

Human Mesangial Cells (HMCs, Innoprot #P10661) were obtained from human healthy kidney. Cells were cultured at 37°C in a humidified environment containing 5% CO_2_, in Roswell Park Memorial Institute (RPMI) medium supplemented with 20% of Fetal Bovine Serum (FBS), 50 U/mL penicillin and 50 µg/mL streptomycin. Cells were used for experiments between passages 4 to 10. HMCs were serum deprived for 24 hours for RT-qPCR and 48 hours (medium replaced at 48, 42, 24, and 2 hours) for western blot before stimulation using RPMI supplemented with 0.5% FBS and antibiotics. Stimulations were done in RPMI 0% FBS and antibiotics, using cells cultured in 6 cells plate during different times for mRNA and protein assays. Albumin (25 µg/mL), β-synuclein (15 µg/mL) (21) (control recombinant protein) and HumanKine® recombinant Platelet-Derived Growth Factor bb (#HZ-1308, 50 ng/mL) were used as negative and positive controls, respectively. Recombinant human soluble CD89 (rsCD89) was produced by the biochemistry and biophysics facility, Inserm U1266, Paris, France, and used at different concentrations. Everolimus (10µM) was used to block mTOR pathway in HMCs one hour before stimulation.

### Western blot analysis

Cells were lysed in RIPA buffer (89900, Thermo Scientific™) supplemented with protease (cOmplete™, Roche) and phosphatase inhibitors (P0044, Sigma-Aldrich). Proteins were denatured in Laemmli buffer (BIO-RAD) with 2.5% 2-Mercaptoethanol and 20–30 μg loaded onto 4%-12% SDS-PAGE gels. Proteins were transferred to Low Fluorescence PVDF membranes (BIO-RAD) and detected using the following primary antibodies: mouse monoclonal anti human-TfR1 (ab269513, Abcam, 1:5,000), mouse monoclonal anti-phospho Akt (Ser473, 66444-1-Ign, Proteintech, 1:5,000), rabbit polyclonal anti-Akt (9272, Cell Signaling, 1:1,000), mouse monoclonal anti-phospho p70S6K1 (Thr389, 9206, Cell Signaling, 1:1,000), rabbit monoclonal anti-p70S6K1 (2708, Cell Signaling, 1:1,000), rabbit monoclonal anti-phospho mTOR (Ser2448, 5536, Cell Signaling, 1:1,000), and mouse monoclonal anti-mTOR (4517, Cell Signaling, 1:1,000). GAPDH was used as a protein loading control with a mouse monoclonal anti-GAPDH antibody (AM4300, Invitrogen, 1:15,000). HRP-conjugated secondary antibodies (Invitrogen) included goat anti-mouse IgG HRP (31430, Invitrogen) and goat anti-rabbit IgG HRP (31460, Invitrogen). Signals were detected using ECL kits (Thermo Scientific™, BIO-RAD) and visualized with the ChemiDoc MP System (BIO-RAD).

### Immunohistochemistry. TfR1 in cIgAN and patients control

Immunostaining was performed on formalin-fixed, paraffin-embedded section using a mouse anti-human TfR1 (ab269513, Abcam, 1:4,000), followed by a polyclonal anti-Ig plus HRP-DAB Novolink polymer detection system (Leica). Tissue sections were read with a laser-scanning microscope (Hamamatsu; nano Zoomer S60).

### Reverse transcription and quantitative polymerase chain reaction (RT-qPCR)

After cells were treated with different stimulants, mRNA was extracted from cells using TRIZOL® Reagent (Invitrogen Life Technologies, Carlsbad, CA, USA). cDNA was obtained from 2,5 µg of total RNA, using Reverse Transcription kit (All-in-one 5X RT MatserMix, ABM). Quantitative real-time PCR analysis of the different genes studied was done using the specific primers, 100 ng of cDNA and following manufacturer’s instructions, with a melting temperature at 60°C, previously determined by gradient PCR (BlasTaq™ 2X qPCR MasterMix, ABM). Relative RNA levels were calculated by normalizing to GAPDH mRNA using 2^−ΔΔCt^ method.

### Single cells analysis

Using data of scRNAseq based on IgAN public dataset (37) (4 IgAN patients and 3 controls) by bioinformatic analysis, we identified and compared subclusters of cells from cIgAN patients and control biopsies to identify complete transcriptomic signature. Using CellPhoneDB, we identified the different cell types and conduct a cell-cell interaction analysis based on the identification of ligand-receptor co-expression (38). We particularly analyzed the expression of mTOR and TfR1 in different types of cells.

### Statistical analysis

Normality was assessed with the Shapiro-Wilk test, while differences between groups were evaluated using the Mann-Whitney and Fisher exact tests, as appropriate. For analyses involving multiple comparisons, either analysis of variance or the Kruskal-Wallis test was applied. Outliers were detected and removed based on the Dixon test. Correlations were measured using the Spearman rank correlation coefficient. All tests applied were two-sided, with a P-value threshold of less than 0.05 for statistical significance. *: P ≤ 0.05, **: P ≤ 0.01, ***: P ≤ 0.001****: P ≤ 0. 0001.Statistical computations were carried out using GraphPad Prism, version 10.

## Results

### TfR1 and mTOR mRNA are overexpressed in mesangial cells from IgAN patients

Single cell analysis of kidneys in IgAN public dataset (37) (4 IgAN patients and 3 controls patients) revealed an increase of mesangial cell numbers in IgAN patients (**Fig. 1A**). This cell subtype showed major increases in TfR1 and mTOR expression as observed in RNAseq analysis **(Fig. 1B)**. Moreover, we detected TfR1 overexpression in the glomerular area by immunostaining in three cIgAN kidney biopsies. These biopsies displayed mesangial expansion, endocapillary hypercellularity, and cellular crescent, whereas the control kidney biopsy was negative for TfR1 staining in the glomerular area (**Fig. 1C)**.

**Figure 1.**
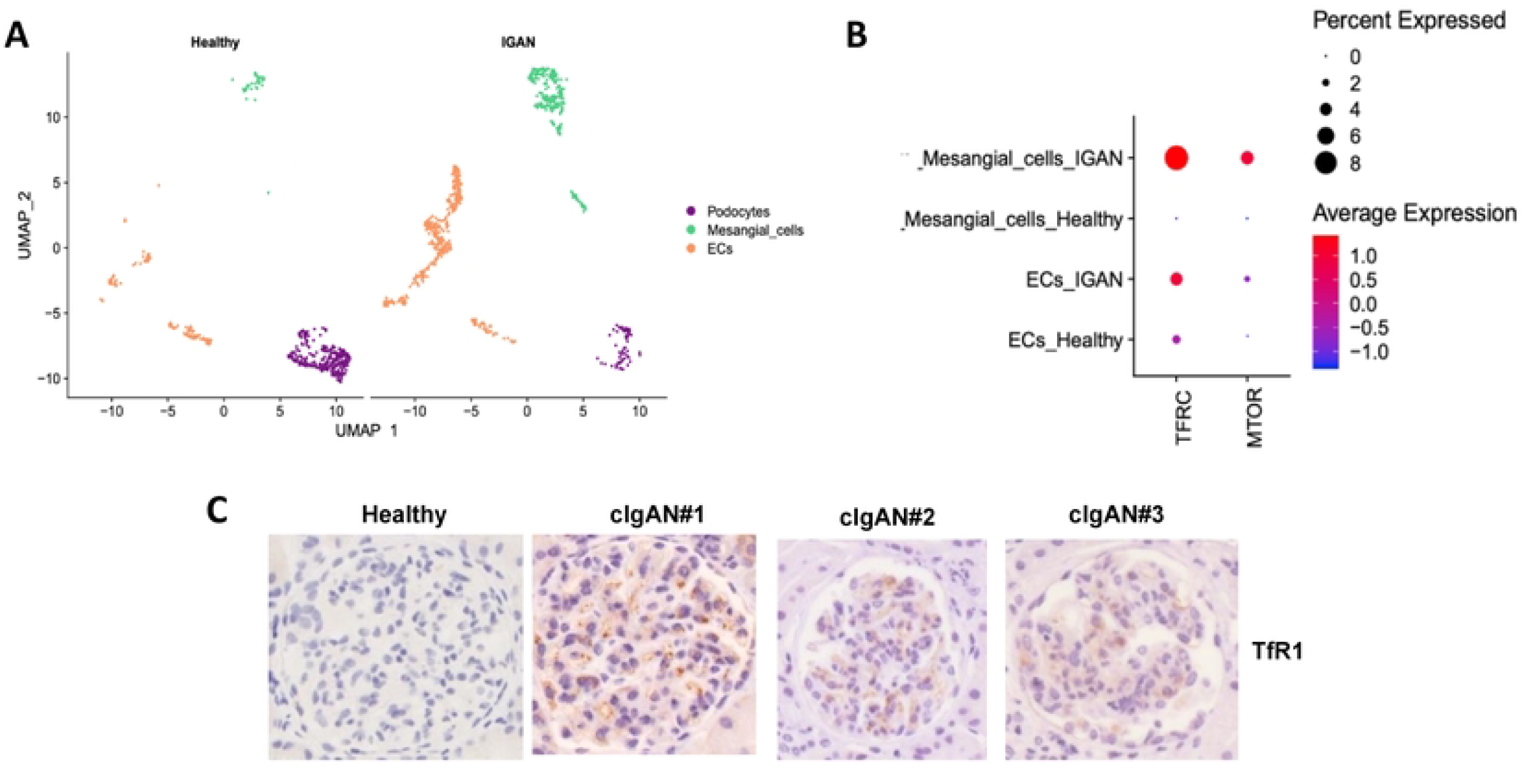
TfR1 and mTOR are overexpressed in human mesangial cells from IgAN patients during single cell analysis and in kidney biopsies of childhood IgAN (cIgAN). (A) Distribution of cell types in the glomeruli of healthy and IgAN patients. Endocapillary cells (Ecs). (B) Dot plot showing gene expression in luminal mesangial cells of TfR1 receptor and mTOR after scRNAseq analysis. (C) Mesangial TfR1 is overexpressed in kidneys from cIgAN patients as compared to those in healthy control patients. Data obtained by immunohistochemistry in childhood biopsies.

### rsCD89 induces mTOR, Akt and p70S6K1 phosphorylation in human mesangial cells

To examine whether sCD89-CD71 interaction induces mTOR activation in mesangial cells, we performed experiments with rsCD89 stimulation of HMCs in vitro in absence of IgA at different time points (20 min, 35 min and 1 hour). rsCD89 (15μg/ml) directly increased expression of TfR1, mTOR, Akt and transglutaminase type 2 (TgM2) at mRNA level **(Fig. 2A)**. At protein level, rsCD89 induced increase in TfR1 expression (**Figure 2B**) followed by phosphorylation changes of mTOR (mTOR p-S2448/mTOR) **(Fig. 2C and F left panels)**, but also of Akt (p-S473 Akt) and S6K1 (p70S6K1) kinases (**Fig. 2D and E**). Everolimus treatment of HMCs for 1 hour resulted in marked decrease mTOR phosphorylation at S2448 (mTOR p-S2448/mTOR), even in the presence of rsCD89 or PDGF **(Fig. 2C and 2F right panel)**.

**Figure 2.**
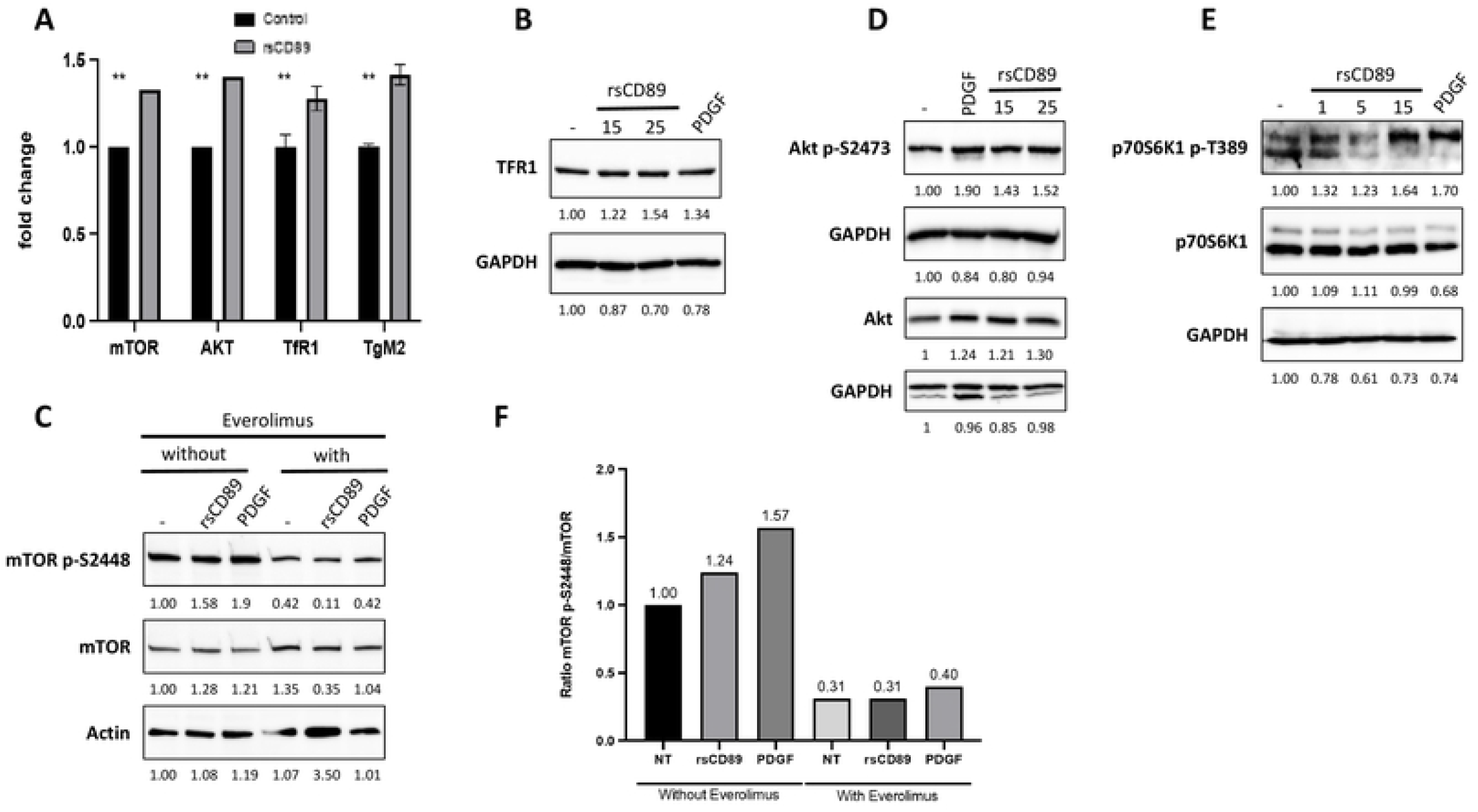
Recombinant soluble CD89 (rsCD89) modulates mTOR and Akt pathways through TfR1 expression in human mesangial cells (HMCs) (A) Quiescent human mesangial cells (HMCs) were stimulated with rsCD89 (15 μg/ml), with β-synuclein used as a negative control. rsCD89 induced the expression of mTOR, Akt, TfR1, and transglutaminase type 2 (TgM2) as measured by RT-qPCR. (B) TfR1 protein expression was upregulated by rsCD89. (C) rsCD89 stimulation induces phosphorylation of mTOR at S2448 (mTOR p-S2448/mTOR). (D) rsCD89 stimulation induced phosphorylation of downstream Akt at S473 (E) rsCD89 stimulation induced phosphorylation of downstream S6K1 kinases at T389. (F) Everolimus treatment of HMCs for 1 hour decreased mTOR phosphorylation at S2448, even in the presence of rsCD89 or PDGF.

### Everolimus prevents IgAN development in the α1^KI^ mice injected by rsCD89

To explore whether mTOR inhibitors could block sCD89-mediated effects in mesangial cells in vivo, we first choose a shorter therapy in the α1^KI^ mice injected with rsCD89 that is able to induce the IgAN disease, as described previously(21, 31). Six α1^KI^ mice received 200 μg rsCD89 twice a week over four weeks, in which with three of them received everolimus (2mg/kg) and three receiving water as a control (**Fig. 3A**). Prior to treatment, IgA1 deposits, proteinuria (microalbuminuria (μAlb)) and cystatin levels were comparable between the groups. After 25 days (D25) everolimus decreased glomerular IgA1 deposits and mesangial sCD89 deposits (**Fig. 3B and C, and Supplementary Fig. 1**), reduced proteinuria levels (**Fig. 3D**), while renal function remained stable as evaluated by cystatin C levels whereas renal function deteriored in the placebo group (**Fig. 3E**). Everolimus decreased also Ki67 staining in kidney (**Fig. 3F)**. The levels of Gd-IgA1 were not affected by the treatment (**Supplementary Fig. 2A**).

**Figure 3.**
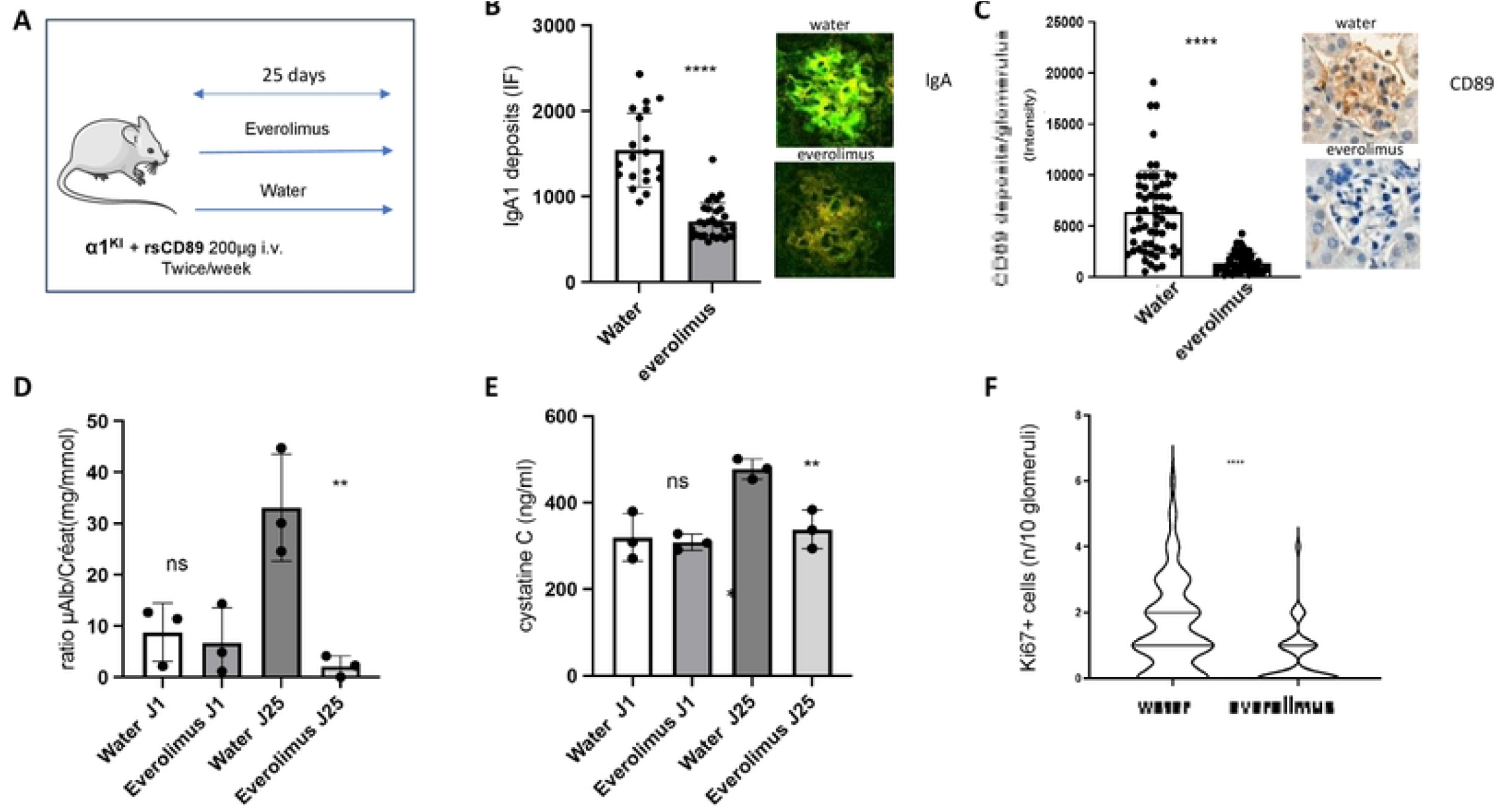
Everolimus prevents IgAN development in the α1^KI^ mice injected by rsCD89. (A). Experimental setup using α1^KI^ mice injected with rsCD89 twice a week. These mice were used to test everolimus as an effective treatment with mTOR inhibitors (2mg/kg/day;) compared to vehicle for 25 days. Quantification of immunostainings were performed in 10 glomeruli per mouse studied as follows: (B). Glomerular IgA1 deposits in immunofluorescence (IF). (C). Glomerular sCD89 deposits in immunohistochemistry. (D). Proteinuria (ratio microalbuminuria (μAlb) /creatinuria mg/mmol). (E). Serum cystatin C levels (ng/ml). (F). Ki67 immunostaining in kidney mice. Representative immunostaining data from water- and everolimus-treated mice are show on the right side of each panel for IgA1 (green) (B) and sCD89 (brown) (D) (mice # 1 and #4, respectively).

### Everolimus reverts IgAN phenotype and halts disease progression in the α1^KI^CD89^Tg^ mice

To validate the mTOR inhibitor effect in mice spontaneously developing IgAN-like disease, we took advantage of the α1^KI^CD89^Tg^ mouse model that progress with time by developping mesangial expansion and IgA1 and sCD89 deposits associated with proteinuria and hematuria (31). To treat initial IgAN model (that might mimic cIgAN disease), ten 8 w-old α1^KI^CD89^Tg^(31) mice were treated daily with everolimus by intragastric gavage for 75 days, while other ten mice received water as a vehicle (**Fig. 4A**). At the endpoint all water-treated animals had a progressive disease as previously described (31). By contrast, everolimus markedly decreased IgA1 deposits in glomeruli (**Fig. 4B and Supplementary Fig. 3**) and sCD89 (**Fig. 4 C and Supplementary Fig. 4)** as well as mesangial TfR1 expression (**Fig. 4 D and Supplementary Fig. 5)**.. Moreover, everolimus-treated group exhibited a significant reduction in C3 deposits, proteinuria (microalbuminuria(μAlb)) and serum cystatine C (**Fig. 4 E**,**F**,**G respectively)**, which were associated with a decrease expression of serum levels of sCD89-IgA1 and IgG-IgA1 complexes (**Fig. 4H and J)**. Gd-IgA1 levels did not reach significance (**Suppl Fig. 2B**). Everolimus induced decreased kidney hypercellularity as quantified by ki67 immunostaining **(Fig. 4K)**.

**Figure 4.**
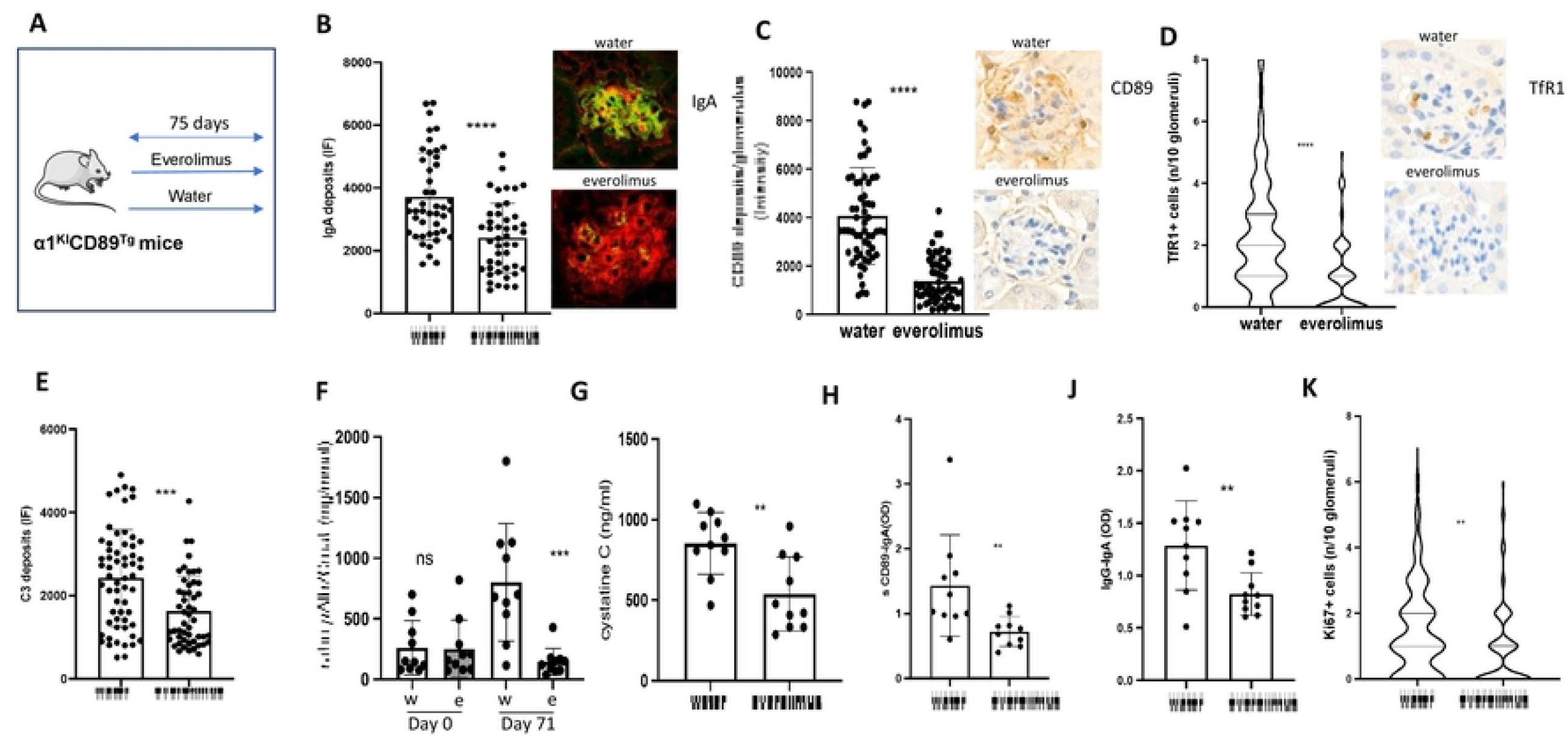
In vivo inhibition of mTOR pathway reverses IgAN progression in the α1^KI^CD89^Tg^ mice. (A) 8-week-old α1^KI^CD89^Tg^ mice expressing human IgA1 and CD89 were used for testing everolimus as effective treatment with mTOR inhibitors (2mg/kg/day; 10 mice) as compared to vehicle (10 mice) for 75 days. Quantification of immunostainings were performed in 10 glomeruli per mouse at the end of the treatment (B) IgA1 deposits, (C) sCD89 deposits, (D) quantification of TfR1 expression and (E) C3 deposits. Representative immunostaining data from water- and everolimus-treated mice are show on the right side of each panel for IgA1(green) (B), sCD89 (brown) (C) and TfR1 (brown) (D) (mice # 1 and #11, respectively). (F) Proteinuria (ratio microalbuminuria (μAlb) /creatinuria mg/mmol) before and after treatment. (G) Serum cystatin C levels at day 75. (H) Serum sCD89-IgA complexes at day 75. (J) Serum IgG-IgA complexes at day 75. (K) QuantificationKi67 staining. Data are shown as mean ± standard error of mean. Statistical significance was determined using a nonparametric Mann-Whitney test or Spearman rank correlation coefficients. (*) p < 0.05, P < 0.05 and ***P < 0.001 compared with the control group. OD, optical density ns (non-significant)

## Discussion

Recent biopharma efforts provided new therapeutic approaches leading to health care progress for patients with IgAN (1, 2, 39). They included mostly B cell and complement modulators but also a spleen tyrosine kinase (SYK) inhibitor (40)). However, despite of promising results in phase 2 and 3 RCT trials, only partial responses were achieved without complete remission. Moreover, no clinical trials exist for these therapies in pediatric populations, limiting treatment options and leaving corticosteroids and renin angiotensin blockers treatment as the primary approach. It remains poorly explored the direct targeting of IgA receptor signaling pathways on mesangial cells to prevent or inactivate the progression of the disease.

IgA receptors are essential effectors involved in IgA-immune responses. Among several IgA receptors, two of them have been shown to be associated with inflammatory diseases (41), notably in IgAN (42). CD89 is an IgA Fc transmembrane receptor expressed on human myeloid cells while CD71 functions as an alternative receptor for IgA1 (41). It is highly expressed on mesangial cells and enterocytes primarily under pathological conditions associated with IgAN, Henoch-Schönlein purpura (HSP), and celiac disease (26, 27, 41). While CD71 binds IgA1, depending on its expression level and on IgA1 glycosylation (29), CD89 can bind both IgA1 and IgA2. However, CD89 soluble form (sCD89) is not commonly found in the circulation of healthy individuals (31). Under healthy conditions, myeloid CD89 controls inflammatory responses notably through interaction with monomeric IgA (mIgA), the second Ig isotype in serum, that induces an anti-inflammatory effect via tyrosine phosphatase recruitment (43, 44). In contrast CD89 binding to polymeric IgA (pIgA) or IgA-containing immune complex aggregates leads to the recruitment of the SYK kinase, triggering pro-inflammatory responses (42, 43).

It has been previously demonstrated that cIgAN patients exhibit two types of IgA1-containing CICs. While large CICs stimulate proliferation of cultured human HMC whereas small complexes inhibit cellular proliferation (45). However, the composition of stimulatory CICs other than Gd-IgA1 has not been investigated notably the role of sCD89. In cIgAN patients, like in adult patients, decreased levels of cell surface CD89 in blood monocytes and neutrophils were observed probably due to a shedding mechanism involving proteases (16, 21) which is associated with the presence of sCD89 in the systemic blood of IgAN patients (14). Interestingly, it has been shown that decrease in sCD89 levels in the blood is associated with severe cases and with IgAN progression towards renal failure in adults (46). This was confirmed in patients with recurrent IgA deposits after renal transplantation (19). In cIgAN, sCD89 binds pathological Gd-IgA1 with higher affinity than physiological mIgA1, promoting pro-inflammatory responses and mesangial IgA1 deposits (47). We recently demonstrated the pivotal role of sCD89 by inducing mesangial cell proliferation in cIgAN. The injection of recombinant sCD89 induced kidney glomerular inflammation and mesangial cell proliferation in transgenic mice expressing human IgA1 (21). Moreover, the levels of sCD89 and sCD89-IgA1 in plasma cIgAN positively correlated with proteinuria, the estimated glomerular filtration rate (eGFR), and with glomerular inflammation M1, E1, C1, and S1 (21). Also, sCD89 and sCD89-IgA1 were more sensitive and specific than proteinuria at predicting glomerular inflammation, endocapillary hypercellularity, and cellular crescents in cIgAN (21).

Similar to CD89, TfR1 (CD71) can also bind polymeric IgA1 and IgA1-containing immune complexes that are able to induce activation of mesangial cells through ERK-PI3K-mTOR signaling pathway leading to receptor overexpression and cell proliferation (21, 30-32). However, CD71 has a preferential binding for Gd-IgA1 rather than normally glycosylated IgA1 that may explain why normally glycosylated monomeric IgA1 cannot induce mesangial receptor alterations in healthy individuals (29).

In the present study, we first took advantage of public data on single cell analysis (37) to show that TfR1 (CD71) mRNA is overexpressed in mesangial cells from four patients with IgAN confirming our previous observations (26, 28). These results were associated with increased in mTOR expression. Immunohistochemistry kidney biopsies of cIgAN patients revealed CD71 overexpression by mesangial cells confirming data previously observed in adult IgAN and in HSP at protein level (26, 28). As sCD89 was previously shown to interact with CD71 mesangial receptor (21, 31), these observations led to us evaluate the effect of sCD89 on mesangial cells in culture. Our results reveal sCD89 directly induces upregulation of CD71 and activation of mTOR pathway. Indeed, phosphorylated adaptors/kinases such as mTOR, Akt and S6K1 were observed following sCD89 triggering of CD71.

mTOR inhibitors are commonly used in clinics (48). A mTOR inhibitor sirolimus has been previously studied in a non-controlled trial as a potential treatment in 23 adult patients with IgAN showing stabilization of renal function, reducing glomerular proliferative lesions. Nevertheless, the interpretation of these results was troubled by its association with another drug, the enalapril (49). Moreover, since 2011 no trial has been performed probably due to the lack of a clear rational for a mechanism of action in IgAN. We evaluated the effect of everolimus using two models of IgAN, the young α1^KI^ mice injected with rsCD89 and the spontaneous IgAN model the α1^KI^CD89^Tg^ mice. In both models, the drug was able either to prevent or to revert established IgAN disease with marked decrease in human IgA1 and sCD89 deposits, mouse C3 deposits, mouse TfR1 (CD71) expression, proteinuria, cystatin C, and the Ki67 proliferation marker. The observed decrease in TfR1 expression by everolimus is particularly intriguing. One can propose a negative feedback mechanism likely driven by the reduction in IgA1 and sCD89 deposits caused by mTOR inhibition. This highlights a compelling feedback loop involving IgA1, CD89, and TfR1. This agrees with previous data showing that the rapamycin effect, another mTOR inhibitor, is able to reduce proteinuria and kidney deposition of rat IgA in an IgAN model developed in rats (50).

## Conclusion

IgA1 and/or sCD89 activate mesangial cells via CD71-mTOR pathway. This study provides a pre-clinical assay for mTOR inhibitor, the everolimus, as attractive therapeutic option in IgAN. It supports novel trials in IgAN, notably in cIgAN where effective treatment targeting mesangial IgA receptors remains poorly provided.

## Acknowledgements

Dr Cyril Cambier from Bichat Hospital provided help in this study. Dr. Gael Cagnone, from the Research Center at Sainte-Justine, contributed his expertise in single-cell analysis.

## Supporting information

**Supplementary Figure 1: Everolimus reduce IgA1 (green) and sCD89 (brown) deposits in the a1KI model after injection of rsCD89**. α1^KI^ mice injected with rsCD89 twice a week. These mice were used to test everolimus as an effective treatment with mTOR inhibitors (2mg/kg/day;) compared to vehicle for 25 days. Glomeruli were stained with anti-human IgA1 antibodies (green) using immunofluorescence and with monoclonal antibody anti-CD89 in immunohistochemistry (brown). # indicates mice numbers

**Supplementary Figure 2: No effect observed on serum Gd-IgA1 levels by everolimus in both IgAN models**.

**Supplementary Figure 3. Everolimus decreased IgA1 deposits in glomeruli of α1**^**KI**^**CD89**^**Tg**^ **mice**. 8-week-old α1^KI^CD89^Tg^ mice expressing human IgA1 and CD89 were used for testing everolimus as effective treatment with mTOR inhibitors (2mg/kg/day; 10 mice) as compared to vehicle (10 mice). Glomeruli were stained with anti-human IgA1 antibodies (green) using immunofluorescence. Representative images of 20 glomeruli (#2–#10 from the vehicle-treated group and #12–#20 from the everolimus-treated group) demonstrate a significant reduction in IgA1 deposits in the glomeruli of mice treated with everolimus compared to those treated with the vehicle. # indicates mice numbers

**Supplementary Figure 4. Everolimus decreased CD89 deposits in α1**^**KI**^**CD89**^**Tg**^ **mice**. 8-week old α1^KI^CD89^Tg^ mice expressing human IgA1 and CD89 were used for testing everolimus as effective treatment with mTOR inhibitors (2mg/kg/day; 10 mice) as compared to vehicle (10 mice). Glomeruli were stained with monoclonal antibody anti-CD89 in immunohistochemistry (brown). Representative images of 20 glomeruli (#2–#10 from the vehicle-treated group and #12–#20 from the everolimus-treated group) demonstrate a significant reduction in CD89 deposits in the glomeruli of mice treated with everolimus compared to those treated with the vehicle. # indicates mice numbers

**Supplementary Figure 5. Everolimus decreased mesangial TfR1 expression in α1**^**KI**^**CD89**^**Tg**^ **mice;** 8-week old α1^KI^CD89^Tg^ mice expressing human IgA1 and CD89 were used for testing everolimus as effective treatment with mTOR inhibitors (2mg/kg/day; 10 mice) as compared to vehicle (10 mice). Glomeruli were stained with monoclonal antibody anti-TfR1 in immunohistochemistry (brown). Representative images of 20 glomeruli (#2–#10 from the vehicle-treated group and #12–#20 from the everolimus-treated group) demonstrate a significant reduction in mesangial TfR1 expression in the glomeruli of mice treated with everolimus compared to those treated with the vehicle. # indicates mice numbers

## Data availability statement

The datasets generated and analyzed during the current study are available from the corresponding author upon reasonable request. Publicly available datasets used in this study, including single-cell RNA sequencing data, can be accessed through (37). All other relevant data supporting the findings of this study are included in the article and its supplementary materials.

## Authors’ contributions

□ **Alexandra Cambier**: Conceptualization, methodology, data acquisition and analysis, writing – original draft, and project supervision.

□ **Jennifer Da Silva**: Performed experiments, data acquisition, and contributed to the writing – review and editing.

□ **Julie Bex**: Conducted experiments and provided critical revisions to the manuscript.

□ **Fanny Canesi**: Data acquisition and analysis, and contributed to the interpretation of findings.

□ **Lison Lachize**: performed experiments, and contributed to data acquisition.

□ **Aurélie Sannier**: Pathology expertise, provided critical insights into histological analyses, and contributed to manuscript revision.

□ **Natacha Patey**: Pathology expertise, provided critical insights into histological analyses, and contributed to manuscript revision.

□ **Erwan Boedec**: Production of soluble proteins and manuscript review.

□ **Amandine Badie**: Contributed to experimental setup, validation, and manuscript preparation.

□ **Renato C. Monteiro**: Conceptualization, funding acquisition, supervision, and manuscript writing and editing.

